# Metformin modulates microbiota-derived inosine and ameliorates methamphetamine-induced anxiety and depression-like withdrawal symptoms in mice

**DOI:** 10.1101/2021.09.30.462054

**Authors:** Jiqing Yang, Zunyue Zhang, Zhenrong Xie, Ling Bai, Pu Xiong, Fengrong Chen, Tailin Zhu, Qingyan Peng, Hongjin Wu, Yong Zhou, Yuru Ma, Yongjin Zhang, Minghui Chen, Jianyuan Gao, Weiwei Tian, Kai Shi, Yan Du, Yong Duan, Huawei Wang, Yu Xu, Yi-Qun Kuang, Juehua Yu, Kunhua Wang

## Abstract

**Objective:** Metformin exhibits therapeutic potential in behavioural deficits induced by methamphetamine (METH) in rats. Emerging studies suggest gut microbiota may impact psychiatric symptoms, but there is no direct evidence supporting metformin’s participation in the pathophysiology of withdrawal symptoms via modulation of gut microbiota.

**Mehods:** In order to define the functional contributions by gut microbiota and metformin to the behavioural deficits during METH withdrawal, we utilized a combination of fecal microbiota transplantation (FMT), high-throughput sequencing, and untargeted metabolomics technologies.

**Results:** First, METH addicts exhibited higher α diversity and distinct microbial structures compared to heathy controls. In particular, the relative abundance of *Rikenellaceae* was positively correlated with the severity of anxiety and depression. Second, both human-to-mouse and mouse-to-mouse FMTs confirmed that METH-altered-microbiota transplantation is sufficient to promote anxiety and depression-like behaviours in recipient germ-free mice, and these behavioural disturbances could be ameliorated by metformin. In-depth analysis revealed that METH significantly altered the bacterial composition and structure as well as relative abundance of several bacterial taxa and metabolites, including *Rikenellaceae* and inosine, respectively, whereas add-on metformin could remodel these alterations. Finally, the inosine complementation successfully restored METH-induced anxiety and depression-like behaviours in mice.

**Discussion:** This study demonstrates that METH withdrawal-induced anxiety and depression-like behaviours are convertible and transmissible via gut microbiota in a mouse model. The therapeutic effects of metformin on psychiatric manifestations are associated with microbiota-derived metabolites, highlighting the role of the gut microbiota in substance use disorders and the pathophysiology of withdrawal symptoms.

**Study Highlights:** *What is known?:*

- There are no targeted therapies for substance withdrawal syndrome, but there is considerable evidence that withdrawal-associated psychiatric manifestations contribute to the poor adherence to rehabilitation treatment as well as the relapse rates.
- Metformin has shown its therapeutic potential against METH-induced neurobehavioural changes and neurodegeneration in rats through CREB/BDNF and Akt/GSK3 signaling pathways in the anxiety-related brain nuclei.

*What is new here?:*

- METH withdrawal-induced anxiety and depression-like behaviours are convertible and transmissible via gut microbiota in a mouse model.
- The therapeutic effects of metformin on psychiatric manifestations are associated with microbiota derived metabolites.
- Inosine complementation could restore METH withdrawal-induced anxiety and depression-like behaviours.

## INTRODUCTION

Methamphetamine (METH) is a potent and long-lasting central nervous system (CNS) stimulant, which is associated with high rates of personal and community harm and remains one of the major public health issues worldwide [1]. Numerous studies have shown that the chronic administration of METH and its abrupt discontinuation cause substance withdrawal syndrome with a series of severe neurobehavioural disturbances including a depressed, anxious, and irritable mood, and difficulty concentrating [2, 3]. Among them, anxiety and depression are the most common psychiatric symptoms emerging during both the METH intoxication and withdrawal stages [4, 5]. Although the prevalence of such psychiatric symptoms has not been well studied in METH-using populations, a few investigations established that over three quarters of chronic METH users [6] and nearly 40% of treatment-seeking METH users [7] reported anxiety and/or depression. To date, there are no targeted therapies for substance withdrawal syndrome, but there is considerable evidence that withdrawal-associated psychiatric manifestations contribute to the poor adherence to rehabilitation treatment as well as the relapse rates.

Mood, cognition, memory, and personality were originally believed to be exclusively modulated by the CNS. However, it is now becoming clear that many extra-neuronal factors, such as the immune system and the gut microbiota that reside in the gastrointestinal tract, could regulate neurological function and have been associated with cognitive neuropsychology and psychosocial functioning [8, 9]. Recently, accumulating clinical and experimental evidence suggest that the alteration of gut microbiota regulates the synthesis of neuroactive molecules and central neurotransmitters, such as γ-aminobutyric acid (GABA), serotonin, dopamine, and melatonin, and therefore may play critical roles in the pathogenesis of anxious and depressive symptoms in neurodegenerative and neuropsychiatric disorders. For example, the gut microflora of patients with Parkinson’s disease contained high levels of *Rikenellaceae* compared to corresponding healthy controls (HCs) and the genera *Turicibacter* and *Prevotella* were significantly correlated with the disease severity scores [10]. The spatial expression pattern of the GABA receptor in the brain could be altered by chronic treatment with *Lactobacillus rhamnosus,* which in turn reduces stress-induced corticosterone levels and depression-like behaviours [11, 12]. In addition, mice that received the fecal microbiota associated with major depressive disorder exhibited depression-like behaviours and disturbances of microbial genes and host metabolites [13]. More recently, it has been reported that the gut-derived isovaleric acid, which is positively correlated with salivary cortisol and depression in boys, could cross the blood–brain barrier and interfere with synaptic neurotransmitter release [14]. Based on these intriguing findings, we hypothesized that the dysbiosis of gut microbiota may participate in the development of psychiatric symptoms in the context of METH addiction and withdrawal via the microbiota-gut-brain axis.

Metformin is a biguanide and it is the most prescribed drug for the treatment of individuals with type 2 diabetes mellitus due to its safety and its glucose-lowering effects [15]. Metformin has also been shown to be beneficial in several other conditions, such as cancer, cardiovascular disease, and neurodegenerative disease [16]. A recent study also showed that metformin could act against METH-induced neurobehavioural changes and neurodegeneration in rats because of its direct activation of the cAMP response element binding protein (CREB)/brain-derived neurotrophic factor (BDNF) and protein kinase B (Akt)/glycogen synthase kinase 3 (GSK3) signalling pathways in the anxiety-related brain nuclei [17]. Mechanistically, metformin is well-tested *in vitro* and *in vivo* and an approved compound that targets diverse pathways including mitochondrial energy production and insulin signaling [18]. In addition, liver, muscle and adipose tissue are classic sites of metformin action, and there is growing evidence from both rodent and human studies suggesting that the gut microbiota might represent another key target involved in the antidiabetic and other possible beneficial effects of metformin [19, 20]. For example, metformin has been proved to modulate gut microbiota composition and structure through increasing mucin-degrading *Akkermansia muciniphila* as well as short chain fatty acid-producing microbiota in patients with diabetes [21]. However, there is no direct evidence supporting that the gut microbiota would be modulated by metformin and become an alternative route participating in the development of substance withdrawal symptoms, and its mechanisms of action remain to be clarified.

In order to explore the effects of metformin on gut microbiota, microbial metabolism, and neurobehavioural symptoms induced by METH exposure, we sought to define functional contributions by metformin and gut microbiota to the behavioural abnormalities associated with METH withdrawal, using a combination of fecal microbiota transplantation (FMT), high-throughput sequencing, and untargeted metabolomics technologies, and most importantly, to pinpoint the underlying interactions and molecular mechanisms of metformin in microbiota-gut-brain axis in the context of METH addiction and withdrawal.

## MATERIALS & METHODS

### Ethics statement and clinical sample collection

Fifteen male methamphetamine addicts (MAs) (age ranging from 18-56) during withdrawal were recruited from the hospital of the sixth Drug Rehabilitation Center in Dehong, China, and 17 age-matched non-substance using controls with no history of any major disease were recruited from the local community. The participants’ age, gender, body weight, and height were collected. Fresh fecal samples were collected from the two groups, frozen immediately, and stored at −80°C for tests. The participant recruitment, fecal sample collection, and clinical information collection and usage was approved by the Ethical Committee from Clinical Research Ethics Committee, the First Affiliated Hospital of Kunming Medical University (2018-L-42). All participants provided written informed consent for sample and clinical data collection and subsequent analyses prior to study participation.

### Scales

The Self-Rating Anxiety Scale (SAS) and the Self-Rating Depression Scale (SDS) were used to measure the level of anxiety and depression for MAs and HCs. The reliability and validity of the Chinese version of these two scales have been confirmed previously [22]. Briefly, both scales contained 20 items and each item was classified as never/rarely, sometimes, often, or always and assigned a score from 1–4, respectively. The testing score was calculated by summing the scores for the 20 items and was standardized by multiplying the sum by 1.25. Scales were measured by experienced a psychological assessor. Higher scores on the SAS or the SDS indicated a higher level of mental disorder [23].

### Animals

Wildtype male C57BL/6 mice, weighting 20-25g were purchased from Hunan Chushang Bioscience Company (Hunan, China). Mice were bred in a specific pathogen-free barrier facility, housed in managed conditions with free access to food and water, and maintained on a 12-hour light/dark cycle (lights on from 07:00 to 19:00) with experimentation occurring during the light cycle. We kept a maximum of five mice per cage in our animal facilities for at least 1 week before use. Research involving mice was approved by the Ethical Committee in the Research Deputy of Kunming Medical University (2020-471) and was performed in accordance with NIH guidelines. All animals were randomly assigned into groups.

### Generation of the germ-free mice

Treatment with a cocktail of broad-spectrum antibiotics is commonly used to deplete the gut microbiota of mice and to generate germ-free (GF) mice. The C57BL/6 mice were administrated with a cocktail of ampicillin (Sigma,1 g/L), metronidazole (Fisher,1 g/L), neomycin (Fisher,1 g/L) and vancomycin (Fisher,0.5 g/L) for 14 consecutive days in drinking water as previously described [24, 25]. Water containing the antibiotics was stored at 4°C before use and changed every 3 days. Mice exhibiting more than a 30% decline in body weight were excluded from the study.

### Fecal microbiota transplantation (FMT)

In the humanized FMT mouse model, donor microbiota were prepared using pooled fecal samples from two MAs with severe depressive symptoms and two age- and gender-matched HCs (figure 1a). In the mouse-to-mouse FMT model, donors were obtained from two mice randomly selected from 10 mice per group. Briefly, 3 days prior to peroral FMT, the antibiotic cocktail was withdrawn and replaced by sterile drinking water. FMT was done as previously described [26]. Fresh fecal pellets from the corresponding donors were immediately weighed and then diluted with sterile PBS (1 g/mL for humanized FMT; 1 fecal pellet/ml for mouse-mouse FMT). The stool was steeped in sterile PBS for about 15 min, shaken, and then centrifuged at 1000 rpm, 4□ for 5 min. The suspension was centrifuged at 8000 rpm, 4□ for 5 min to get total bacteria, then filtered twice in PBS. For each GF recipient mouse, 200 μl of bacterial suspension (10^8^ CFU/mL) was transplanted by gavage each day for 7 consecutive days. The recipient mouse was maintained for 7 days for transplanted microbiota recolonisation before being subjected to experiments.

**Figure 1.**
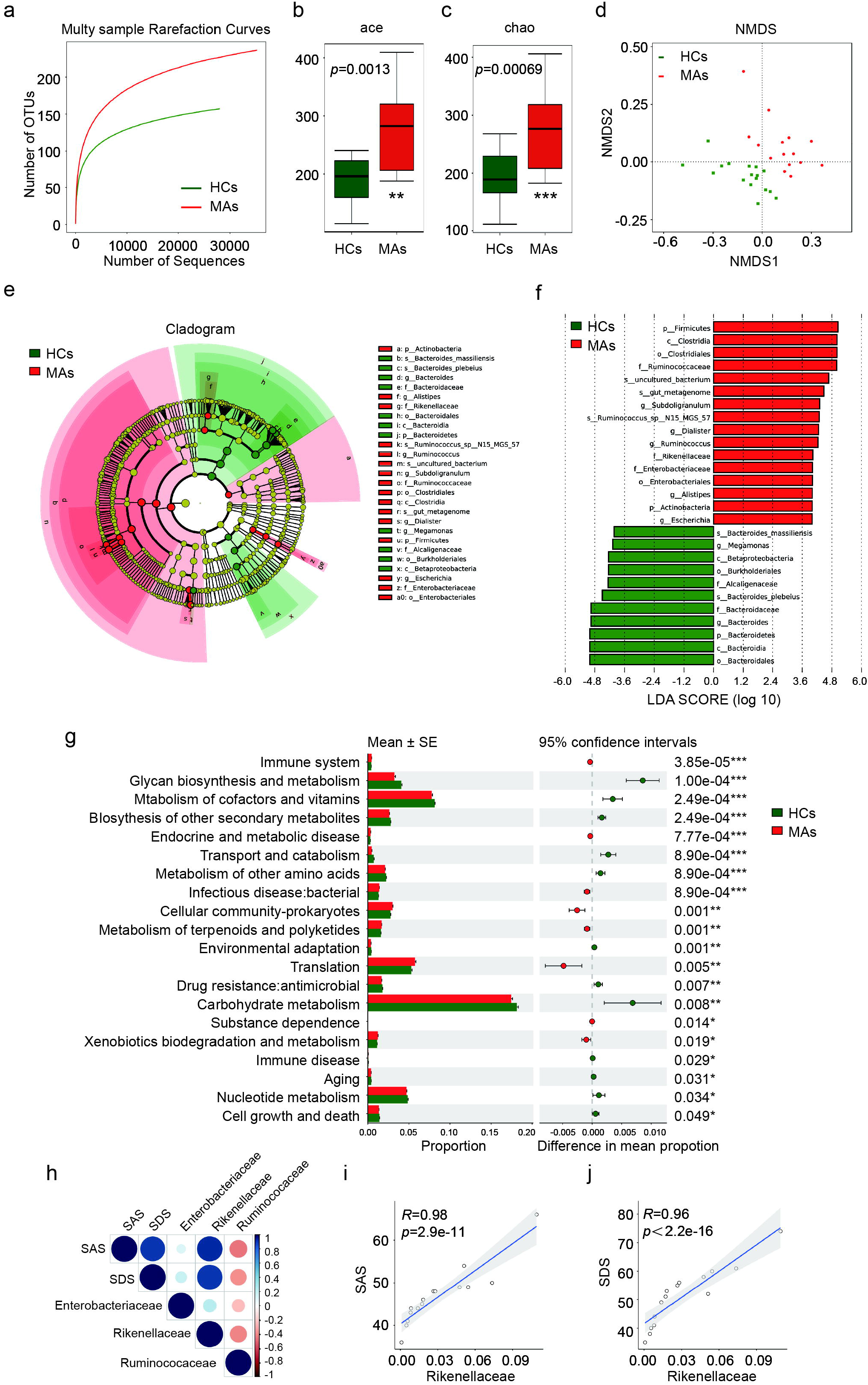
Microbial characteristics of MAs. Alterations in intestinal microbiota diversity, composition, and function in MAs. (a) The observed species rarefaction curve of the two groups. Rarefaction curves were assembled showing the number of OTUs, relative to the number of total sequences indicated that the abundance of species in gut microbiota in MAs is higher than in HCs. (b-c) α diversity index ace and chao of the two groups. A higher diversity index was observed in MAs compared to the HCs according to ace (*p* = 0.0013) and chao (*p* = 0.00069) by the Wilcoxon rank-sum test. (d) Nonmetric multidimensional scale method (NMDS) analysis showed that the gut microbiota composition can distinguish samples from MAs and HCs. (e) Cardiogram showing differentially abundant taxonomic clades with a linear discriminative analysis (LDA) score > 4.0 among MAs (red) and HCs (green), *p* < 0.05. (f) Linear discriminative analysis (LDA) effect size (LEfSe) analysis between the HCs (red) and MAs (green), (g) Differences in the notability function in the Kyoto Encyclopedia of Genes and Genomes (KEGG) module prediction using 16S data with PICRUSt2. (h-j) Correlation between the relative abundance of the families *Rikenellaceae, Ruminococcaceae, Enterococcaceae, Bacteroidaceae, Alcaligenaceae* and SAS and SDS. Each dot represents an individual and correlation was calculated using Pearson’s correlation.

### METH and metformin treatment in mouse model

METH was obtained from Narcotics Department of Yunnan Provincial Public Security Administration and was dissolved in 0.9% NaCl solution. Metformin (Sigma-Aldrich, USA) were dissolved in 0.9% NaCl solution. Mice were randomized to one of four treatment groups: Saline, METH, Metformin and METH/Metformin program. Mice were treated with Saline (0.2ml/mouse, i.p.), METH (5mg/kg, i.p.), Metformin (200mg/kg, i.p.) and METH/Metformin (i.p.) for 21 days on the corresponding treated group respectively. The METH dose was determined based on the data of behavioural sensitization and CPP test (supplementary figure 1) while metformin dose was determined as previously used in preclinical [27] and animal study [28, 29]. Body weights were measured every week, fecal sample were collected on day 24.

### Behavioural tests

All behavioural analyzes were performed during the 09:00–17:00 light cycle. Animals were habituated in the test room for 2-h before starting the experiments. The open field and elevated plus apparatus were cleaned with 70% ethanol between each trial. To establishment of METH dependent mouse model, mice were treated with METH (5 mg/kg, i.p.) once a day for 21 consecutive days (figure 2b). Behavioural procedures locomotor sensitization and conditioned place preference (CPP) which were associated with rewarding effects of METH were used to confirm the establishment of addiction and dependent. The behavioural tests were conducted in this order: open field test (OFT), elevated plus maze (EPM), tail suspension test (TST) and forced swim test (FST). All behavioural analyzes were performed blinded to treatment groups.

**Figure 2.**
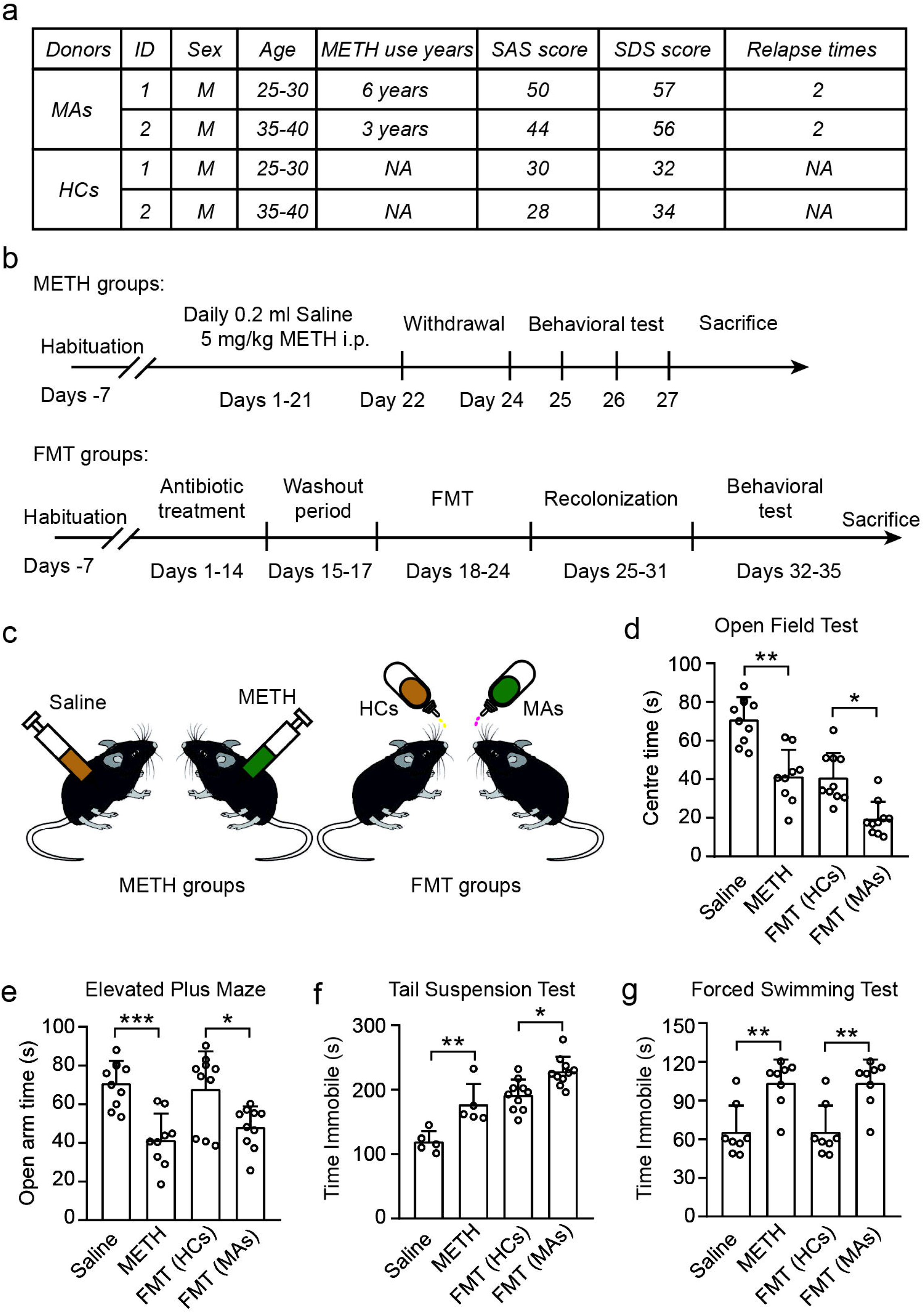
Colonization of GF mice with MA microbiota reproduce human behaviours. Transferring FMT from MAs with high Self-Rating Anxiety Scale (SAS) and Self-Rating Depression Scale (SDS) scores to microbiota-depleted mice induced anxiety and depression-like features in the recipient animals. (a) Metadata of four human donors used for mouse colonization. (b-c) Experimental design. (d-g) Behaviour test. Panel d shows METH withdrawal mice and transplantation of the fecal microbiota from MAs. GF mice spent less time in the centre area in the open field test than control mice. Panel e shows results of time spent in the open arm in the elevated plus maze. Panel f shows results of immobile time in tail suspension test (n=5 in METH group and n=10 in FMT group). Panel G shows results of immobile time in the forced swim test. All data are expressed as mean ± SD.

### Behavioural locomotor sensitization

METH-induced behavioural sensitization was measured in an open field test using the ENV-510 test environment and ANY-maze software (Stoelting Co.) as described in previous study [30]. Mice received an injection of normal saline (10□ml□/kg) as control and 5 mg/kg METH as treated group for 13 consecutive days. On day 14, mice were injected with METH or saline immediately before confinement to the test room, locomotor activity was monitored for 60□min.

### Conditioned place preference (CPP)

METH-induced CPP was evaluated using the CPP system and monitored by ANY-maze (Stoelting Co.). Briefly, the CPP schedule consisted of three phases: preconditioning, conditioning, and post-conditioning. In preconditioning phase, mice were initially placed in the middle chamber with the doors removed for 15 min as the baseline preference. During the conditioning phase, mice was treated for 14 days with alternate injections of either METH (5 mg/kg, i.p.) or saline (2 ml/kg, i.p.). Mice were confined to the white compartment for 45 min immediately after METH administration and to the black compartment after saline injection. In the postconditioning phase, CPP testing was carried out on day 15 when each animal was again allowed to explore all chambers freely, the time spent in each of the two compartments was automatically recorded for 15 min. CPP scores were defined as post-conditioning time subtracted from preconditioning time.

### Open field test (OFT)

To observe subsequent behaviours for evaluating anxiety and locomotor activity, each mouse was placed individually in the corner of an open-field arena (45 ×45 ×30 cm) and allowed to explore freely for 6 min. Its spontaneous activities were recorded using a video tracking system (ANY-maze, Stoelting Co.). The total movement distance was used as a measure of locomotor activity, while the time spent in the center (inner 25% of the surface area) was used as an index of anxiety-like behaviours [31].

### Elevated plus maze (EPM)

EPM was used to determine the unconditioned response to a potentially dangerous environment. Anxiety-related behaviour was measured by the degree to which the rodent avoided the open arms of the maze. As previously described [32], on each of the assessment days, mice were transferred to the middle of the elevated plus maze apparatus, and allowed freely to explore within the four arms for 5 min. The time of activity at open arms (s) were recorded by the video tracking system (ANY-maze, Stoelting Co.) as indicative of anxiety-like behaviour.

### Forced swim test (FST)

The FST evaluates the depressive-like behaviour in rodents. A day before the test, all mice were gently placed in cylinder (30 cm height, 17 cm diameter) filled with water to a depth of 15 cm at 25°C and made to swim for a habituation period of 15 min. However, during experimentation, subjects were placed individually in filled glass cylinder for a period of 6 min and the total duration of immobility was measured (ANY-maze, Stoelting Co.) as indicative of depressive-like behaviour. Immobility was defined as floating or remaining motionless without leaning against the wall of the cylinder [33].

### Tail suspension test (TST)

In TST, mouse was suspended 50 cm above the surface of a table using adhesive tape placed 1 cm away from the tip of the tail, mice were considered immobile only when they hung passively and were completely motionless. We recorded duration of immobility in a 6-minute period by a video tracking system (ANY-maze, Stoelting Co.) which was indicative of depressive-like behaviour [34].

### 16S rRNA sequencing analysis

Microbial genomic DNA was extracted from fecal samples following the manufacturer’s protocol, using the MagPure Stool DNA KF Kit B (Magen, China). The quantity of genomic DNA was verified using the Qubit dsDNA BR Assay kit (Invitrogen, USA). The V4 regions of the 16S rRNA gene in the DNA extracted from fecal samples were amplified using the following degenerate PCR primers:515 F (5’-GTG CCA GCM GCC GCG GTA A-3’) and 806R (5’-GGA CTA CHV GGG TWT CTA AT-3’). 16S rRNA sequencing analysis and its diversity was analyzed was performed using a combination of software mothur (version 1.33.3), UPARSE (usearch version v8.1.1756, http://drive5.com/uparse/), and R (version 3.6.3) as previously described [35]. The represent sequences of OTU were classified with Silva database (version 128) with confidence score ≥ 0.6 by the classify.seqs command in mothur. Rarefaction curves were generated based on OTU. The α-diversity analysis was calculated using mothur. For the β-diversity analysis, nonparametric multi-dimensional scaling (NMDS) plots were depicted using the Vegan package. Discriminant analysis was performed using the linear discriminant analysis (LDA) effect size (LEfSe) pipeline. PICRUSt2 was used to identify differences in the metabolic pathways between each group against the KEGG.

### Untargeted metabolomic relative quantitative analyzes

The LC–MS analysis was performed as described previously [36]: 20 mg of fecal samples were accurately weighed and collected, mixed with adequate amounts of precooled acetonitrile/methanol (1:1, v/v), centrifuged for 20 min at 4 °C and 14,000g to collect the supernatant. Metabolic profiling of fecal samples was performed using an ultra-high-performance liquid chromatography (UHPLC, 1290 Infinity LC, Agilent Technologies, Santa Clara, CA, USA) coupled with a quadrupole time of-flight system (AB Sciex Triple TOF 6600, AB SCIEX) at Shanghai Applied Protein Technology Co., Ltd. The raw mass spectrometry data were converted to MzXML files using Proteo Wizard MSConvert before being imported into freely available XCMS software. Principal component analysis (PCA) and partial least square discriminant analysis (PLS-DA) were performed for both positive and negative models after log transformation and pareto scaling.

### Inosine complementation

To test the effects specific metabolites inosine on behavioural phenotypes, mice were supplemented with 300mg/kg inosine (Sigma-Aldrich, USA) by intraperitoneal injection 4h after METH treatment. The dose was determined in mice based on previous described [37]. Body weight was measured at baseline and post-treatment.

### Statistical analysis

Statistical analysis was performed using Prism software (GraphPad). Data are represented as mean±SD or mean±SEM. The differences between two groups were assessed using a two-tailed, unpaired t-test. The differences among three or more groups were assessed using one-way or two-way ANOVA. Wilcoxon test and Kruskal-Wallis test were used to evaluate differences in the microbiota between two or multiple groups. Correlations between variables were calculated using Spearman’s rank-correlation analysis with R version 3.5.3. Significant differences are indicated in the figures by *p<0.05, **p<0.01, ***p<0.001, **** p<0.0001. Notable nearly significant differences (0.05<p<0.1) are indicated in the figures.

## RESULTS

### The gut microbiota profile is altered in METH addicts

To characterise the gut microbiota from MAs, we enrolled a total of 32 study participants with an average age of 37.75±11.12 years, including 15 male MAs currently undergoing withdrawal and 17 age- and gender-matched HCs. There were no statistically significant differences in demographic or anthropometric parameters between the two groups of study participants. Compared to the HCs, the levels of anxiety (46.87±6.94 vs. 40.58±4.39, *p* = 0.0042) and depression (51.1±10.36 vs. 38.47±8.73, *p* = 0.0008) were significantly higher in the MAs (table 1).

**Table 1.** Demographic and clinical characteristics of the study participants. There were no statistical difference of age and BMI in two groups. MAs were in anxiety and depression during withdrawal according to the scores of SAS and SDS. Data are expressed as mean ± SEM; significance testing is by paired t-test.

|  | HCs | MAs | p |
| --- | --- | --- | --- |
| Subjects | n=17 | n=15 | NA |
| Age (years) | $38.24 \pm 12.13$ | $37.20 \pm 9.99$ | 0.7974 |
| Gender | Male | Male | NA |
| BMI (kg/m<sup>2</sup>) | $22.8 \pm 2.97$ | $22.9 \pm 3.37$ | 0.591 |
| METH use time (y) | NA | 3 |  |
| Relapse times (n) |  |  |  |
| 1st time | NA | 5 |  |
| 2nd time | NA | 9 |  |
| More than twice | NA | 1 |  |
| SAS | $40.58 \pm 4.39$ | $46.87 \pm 6.94$ | **0.0042** |
| SDS | $38.47 \pm 8.73$ | $51.13 \pm 10.36$ | **0.0008** |

In the present gut microbiota investigation, we surveyed bacterial composition by 16S rRNA gene deep sequencing and generated over 1,476,500 (~ 450 bp) raw sequencing data. After demultiplexing and quality filtering, we obtained a total of 1,325,000 high-quality reads (mean, 40,000 reads/sample) and first evaluated the ecological features of the bacterial communities between the two groups using a variety of indices. Based on the rarefaction analysis estimates, the species richness in MAs was much higher than that of HCs (figure 1a). Richness estimates such as ace and chao1 indices also indicated that the bacterial α diversity was significantly higher in the MAs than that in the HCs (figure 1b-c). Simultaneously, the nonmetric multidimensional scaling (NMDS) analysis for β diversity revealed a distinct structural difference between the two groups (figure 1d). Taken together, the diversity analyses indicated that the gut microbiota from MAs with relatively higher α diversity and distinct microbial structures was significantly different than those from HCs. Beyond the general composition of the microbiota, specific taxa were also observed to be differentially expressed between MAs vs HCs. Linear discriminant analysis (LDA) identified five statistically significant differences between the two groups at the family level (LDA> 4.0, p<0.05). The relative proportions of *Ruminococcaceae, Rikenellaceae,* and *Enterobacteriaceae* were significantly higher in MAs compared with HCs. We also found significantly lower levels of *Bacteroidaceae and Alcaligenaceae* in MAs than in HCs (figure 1e-f), suggesting that these differential bacteria could be considered as potential biomarkers. In addition, the functional diversity of the putative metagenomes was assessed using the PICRUSt, allowing the prediction of signalling pathways from the 16S rRNA data. As shown in figure 1g, there were significant differences in the mean proportions between the two groups, and some pathways displayed a difference of at least 0.1%. Specifically, the pathways including immune system (p=3.85e-05), endocrine and metabolic disease (p=7.77e-04), and bacterial infection (p=8.90e-04) were significantly enriched in MAs, suggesting that the gut microbial alterations in MAs may be involved in endocrine/metabolic and infectious diseases.

Furthermore, we evaluated correlations between the relative abundances of bacterial families and the severity of withdrawal symptoms using the Spearman correlation method. With significant inter-individual variability, we identified that the relative abundance of *Rikenellaceae* was positively correlated with both SAS (p<0.001, r=0.98) and SDS (p<0.001, r=0.96) scores (figure 1h-j). These data suggest that the higher level of relative abundance of *Rikenellaceae* might predict more severe anxious and depressive symptoms in MAs.

### Gut microbiota from MAs with anxiety and depression is sufficient to promote behavioural deficits in mice

Evidence exists that the gut dysbiosis is associated with METH withdrawal induced depressive behaviours in rats [38]. We sought to determine whether the transplantation of human gut microbiota was sufficient to transfer the hallmarks of the withdrawal syndrome state from MAs to GF mice. Herein, we utilized a practical and clinically relevant GF mouse model by treating mice with an antibiotic cocktail (MATERIALS AND METHODS), after which the depletion of gut microbiota was confirmed (online supplementary figure 2). Two representative MA donors were chosen based on their SAS/SDS scores and history of substance use and relapse. Consistent with previous analysis, the fecal samples from MAs exhibited significant alterations in both α and β diversities within bacterial communities compared to the HCs donors (data not shown). After that, the fecal samples from MAs and HCs donors were transplanted into GF mice to generate “humanized microbiota” mice, denoted as FMT-MAs and FMT-HCs, respectively.

METH-dependent mice were also generated (online supplemental figure 1a). Consistent with the psychiatry symptoms frequently observed in MAs, these mice exhibit severe behavioural deficits during the acute withdrawal stage (online supplemental figure 1b-c & figure 2a). Similarly, these FMT-MAs mice exhibited significantly decreased central time in the open field test (OFT) and the open arm time in the elevated plus maze (EPM) test when compared to the FMT-HCs mice (figure 2d-e). In addition, these FMT-MAs mice showed increased immobility time in the tail suspension test (TST) and forced swim test (FST) tests compared to the FMT-HCs mice (figure 2f, g). Overall, these results indicated that the FMT*-*MAs mice displayed obvious anxiety and depression-like behaviours, suggesting that the FMT transfer from human MA donors into GF mice could also transfer withdrawal syndrome-relevant behavioural deficits.

### Administration of metformin ameliorates METH-induced anxiety and depression-like behaviours in mice

Metformin has been reported to ameliorate METH-induced depression-like neurobehaviours in rats [17]. To investigate the underlying molecular mechanism regarding metformin’s action on behavioural deficits associated with METH withdrawal, mice were grouped and treated with saline, METH, metformin, and METH/metformin (figure 3a). In accord with previous results, mice spent much less time in the central square of the OFT and spent longer time in open arms but less time in closed arms in the EPM after a 21-day METH exposure and then an abrupt cessation, indicating that these mice in the METH group displayed a relatively more severe anxious-like behaviour compared to the mice in the saline and metformin groups. There were no obvious behavioural changes in the metformin group mice in either OFT nor in EPM tests (figure 3b-d). It is worth noting that although metformin did not ameliorate METH-induced hyperactivity in OFT (figure 3c), it could largely reverse METH withdrawal-induced anxiety in mice. For example, mice treated with METH/metformin spent a significantly prolonged time in the central region of the OFT but decreased time in the closed arms of the EPM as compared to the mice treated with METH only. We further performed FST and TST to assess depression-related behaviours in mice. METH significantly prolonged the immobility time of mice in the TST and shortened the swim time but prolonged immobility time in the FST when compared to the mice in saline or metformin groups. Notably, the immobility could be abolished by adding metformin treatment, and results were statistically significant in comparison to mice receiving METH administration only (figure 3e-f). All these results provide additional evidence that metformin ameliorates METH withdrawal-induced anxiety and depression-like behavioural disturbances.

**Figure 3.**
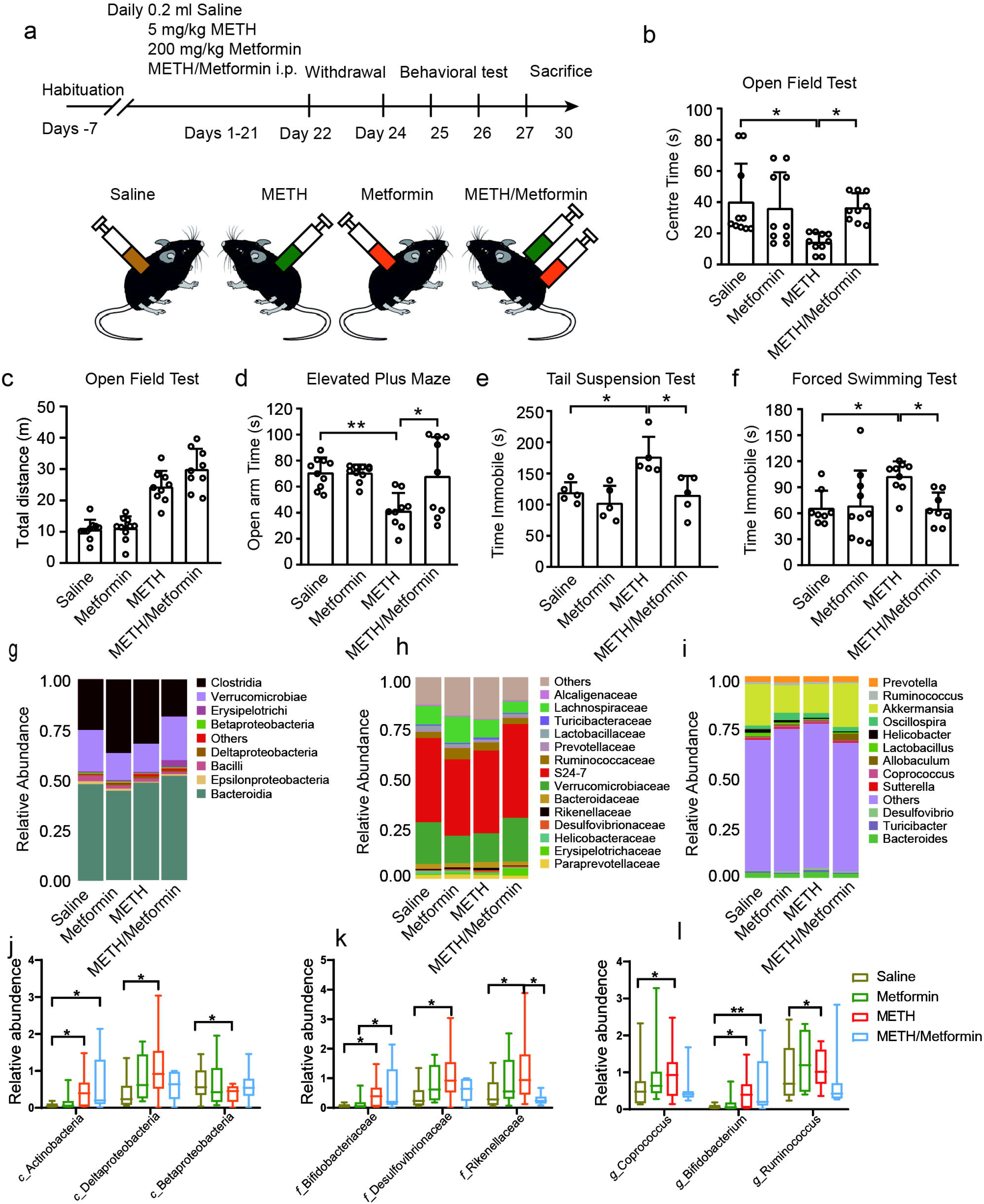
Metformin improved METH withdrawal-induced anxiety and depression behaviour and characteristics of gut microbiota in all groups. (a) Experimental scheme. (b-c) Results of the open field test in all groups. (d) Results of the elevated plus maze in all groups. (e) Results of the tail suspension test in all groups. (f) Results of the forced swimming test in all groups. (g-i) Taxonomic distribution at the class (g), family (h) and genus (i) level in the four groups. Different colours indicate different flora, and the length of each colour column represents the abundance of corresponding flora. Abundance is presented in terms of the percentage of the total effective bacterial sequences in each group (n=10). (j-k) Statistically significant differences taxa are listed (p value of Wilcox test <0.05) at the class (j), family (k) and genus (l) level.

To further determine whether the metformin could modulate METH-altered gut microbiota and therefore participate in the development of withdrawal symptoms, we investigated their fecal microbiota profiles from the four above mouse groups and analyzed the alterations of microbial composition at the class, family, and genera levels (figure 3g-i). At the class level, the relative abundances of *Actinobacteria* and *Detaproteobacteria* in the METH group was higher than in the saline group, whereas the relative abundances were lower in METH/Metformin-treated mice compared to the METH group. Meanwhile, the relative abundance of *Beltaproteobacteria* in the METH group mice was higher than that in the saline group mice, whereas they were less abundant in the METH/Metformin group mice compared to the METH group. At the family level, the relative abundance of *Bifidobacteriaceae, Desulfovibrionaceae,* and *Rikenellaceae* were significantly increased in METH group mice compared to the saline group mice. The METH/Metformin-treated mice had a higher relative abundance of *Bifidobacteriaceae,* whereas the relative abundance of *Desulfovibrionaceae* and *Rikenellaceae* were significantly decreased (figure 3h, k). At the genus level, the relative abundance of *Coprococcus, Bifidobacterium,* and *Ruminococcus* in the METH-treated mice was increased compared with control mice, whereas metformin decreased the relative abundances of *Coprococcus* and *Ruminococcus* but the decrease was not significant (figure 3i, l). Overall, METH exposure significantly altered the bacterial composition and structure, as well as relative abundance of a number of bacterial groups, whereas the added metformin treatment partially reversed these alterations.

### Administration of metformin restores METH induced microbial disturbances that correlate with behaviours

To determine whether the metformin-altered microbiota contribute to the modulation of METH withdrawal syndrome-relevant behavioural deficits, we further conducted mouse-to-mouse FMT by transferring fecal samples from three groups of treated mice (METH, saline, and METH/metformin) in the GF mouse model (figure 4a). After 1 week of colonisation, the “METH microbiota” recipient GF mice displayed a decreased centre time in the OFT and spent less time in the open arm in the EPM compared to “saline microbiota” recipient GF mice, which is indicative of anxiety-like behaviours in METH microbiota recipient mice (figure 4b-d). In the TST and FST, the immobility time in METH microbiota recipients was significantly longer than that of controls in saline group, indicating a stronger depression-like phenotype in these mice (figure 4e-f). However, the METH/metformin microbiota recipient mice continued to exhibit less severe depressive and anxiety behaviours compared to the METH microbiota recipient mice (figure 4b-f). These results indicated that the METH-induced anxiety and depression-like behaviours and the reversal effect of metformin on the withdrawal symptoms were transmissible via the gut microbiota.

**Figure 4.**
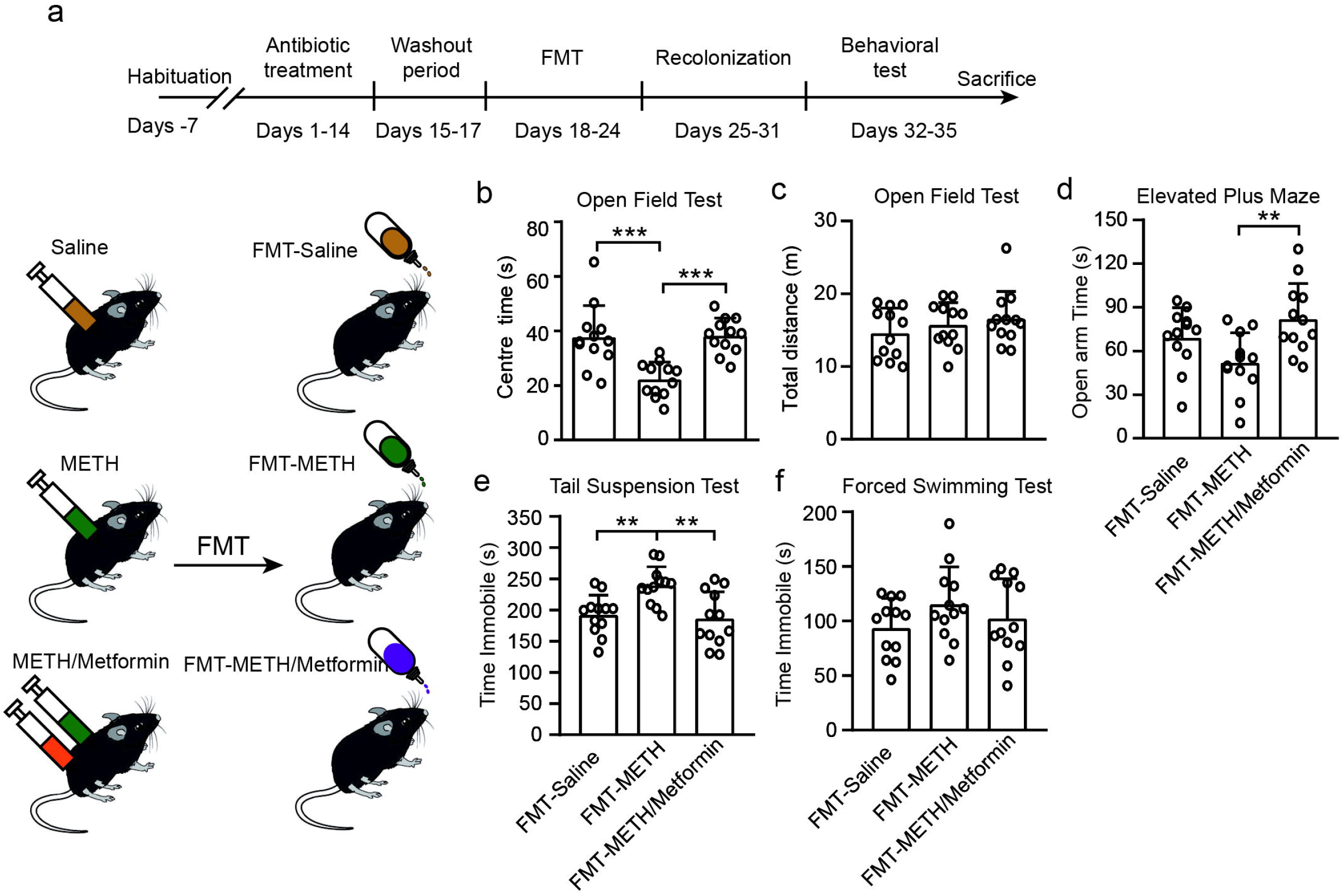
Transfer of fecal samples from mice-to-mice showed that withdrawal symptoms were improved in mice that received metformin-altered microbiota. (a) Experimental scheme. (b-f) Mice receiving FMT from METH-treatment mice showed the same behaviour as donor mice; mice transplanted with metformin-treated fecal microbiota showed lower anxiety-like and depression-related behaviours. (b-c) Time in the centre zone and locomotor activities in open field test were analyzed. (d) Time spent in the open arm in the elevated plus maze test. (e) Time spent immobile in the forced swimming test. (f) Time spent immobile in the tail suspension test. All data are expressed as mean ± SD (n=12/group).

To identify the bacteria that might be responsible for the effects exerted by FMT, the composition of the gut microbiota in caecal content after FMT was further analyzed. There were significant differences at family level, which were composed by *Verrucomicrobiaceae, Rikenellaceae, Erysipelotrichaceae, Helicobacteraceae, Prevotellaceae, Bacteroidaceae, Porphyromonadaceae, Peptostreptococcaceae,* and *Lachnospiraceae.* Of these, *Verrucomicrobiaceae* and *Rikenellaceae* were significantly different at the family level between three groups as determined by LEfSe (online supplementary figure 4). Taken together, our findings support the notion that altered gut microbiota mediate metformin’s anti-anxiety and anti-depression effects.

### Untargeted metabolomics revealed an association between action of metformin and microbiota-derived metabolites

Metabolites of commensal bacterial play a key role in microbe–host interactions [39, 40]. Metabolomic analyzes of fecal samples were also performed in FMT mouse models to determine the bacterial metabolite changes in response to METH withdrawal and metformin treatment using liquid chromatography-mass spectrometry. A total of 357 volatile organic compounds were identified in our untargeted metabolomics analysis from 54 fecal samples in all six groups. Subject-specific compounds and metabolites present in less than 20% of subjects in both groups were discarded from statistical analysis. To identify differences in metabolic profiles among saline, METH, and METH/Metformin-treated groups, as well as in the FMT group, PLS-DA score plots were performed for both donor and recipient modes. The PLS-DA score plot from the fecal samples (R 2X=0.0.188, R 2Y=0.793, Q2=0.337 in the donor groups and R 2=0.0.487, R 2Y=0.876, Q2=0.7 in the recipient groups) illustrates excellent metabolic distinctions among saline, METH, and METH/metformin groups and in the recipient groups (figure 5a). In total, 13 metabolites showed significant group differences (figure 5b), of which inosine, deoxyinosine, folinic acid, ketoisocaproic acid and allantoin were significantly decreased or almost totally absent in faeces from METH-treated mice but were nearly completely restored after metformin treatment (figure 5c-h).

**Figure 5.**
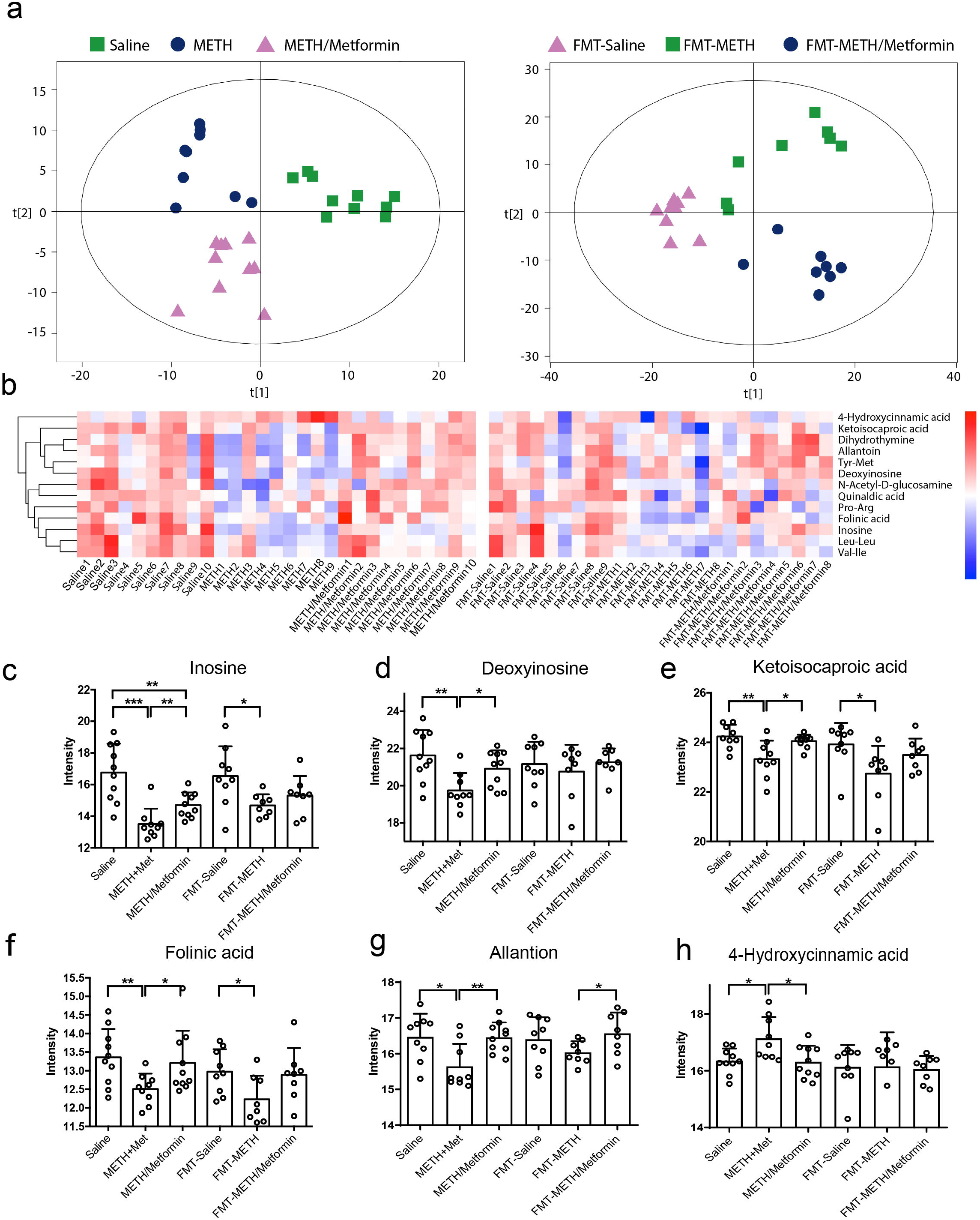
Fecal metabolomic profiles are modulated by metformin treatment. (a) Scatter plot of PLS-DA scores for fecal metabolites showing significant separation among saline, METH and METH/metformin groups as well as in the recipient mice. (b) Heat map showing the levels of 13 fecal metabolites that were significantly altered by metformin treatment in METH-dependent mice. Colours indicate fold changes with *p* < 0.05. (c-h) Relative quantification of metabolites related in fecal samples (one-way ANOVA; n=8~10/group). Error bars represent mean ± SEM.

### Inosine complementation normalizes METH induced anxiety and depression-like behaviours in mice

Inosine is a common component of food and has been shown to have a potential neuroprotective function [41]. To test the hypothesis whether inosine complementation could be beneficial for substance withdrawal syndrome, METH-treated mice were given inosine (300 mg/kg, i.p.) for 3 weeks and then underwent behavioural testing (figure 6a). Inosine complementation significantly increased the time mice spent in the centre in the open field test and the duration in the open arms of the elevated plus maze as compared to the METH group mice (figure 6b-d). In addition, mice treated with METH/inosine treatment had significantly reduced immobility time in the tail suspension test and the forced swim test compared to the mice in the control group (figure 6e, f). Combined, these data suggest that inosine complementation could restore, at least partially, METH withdrawal-induced anxiety and depression-like behaviours.

**Figure 6.**
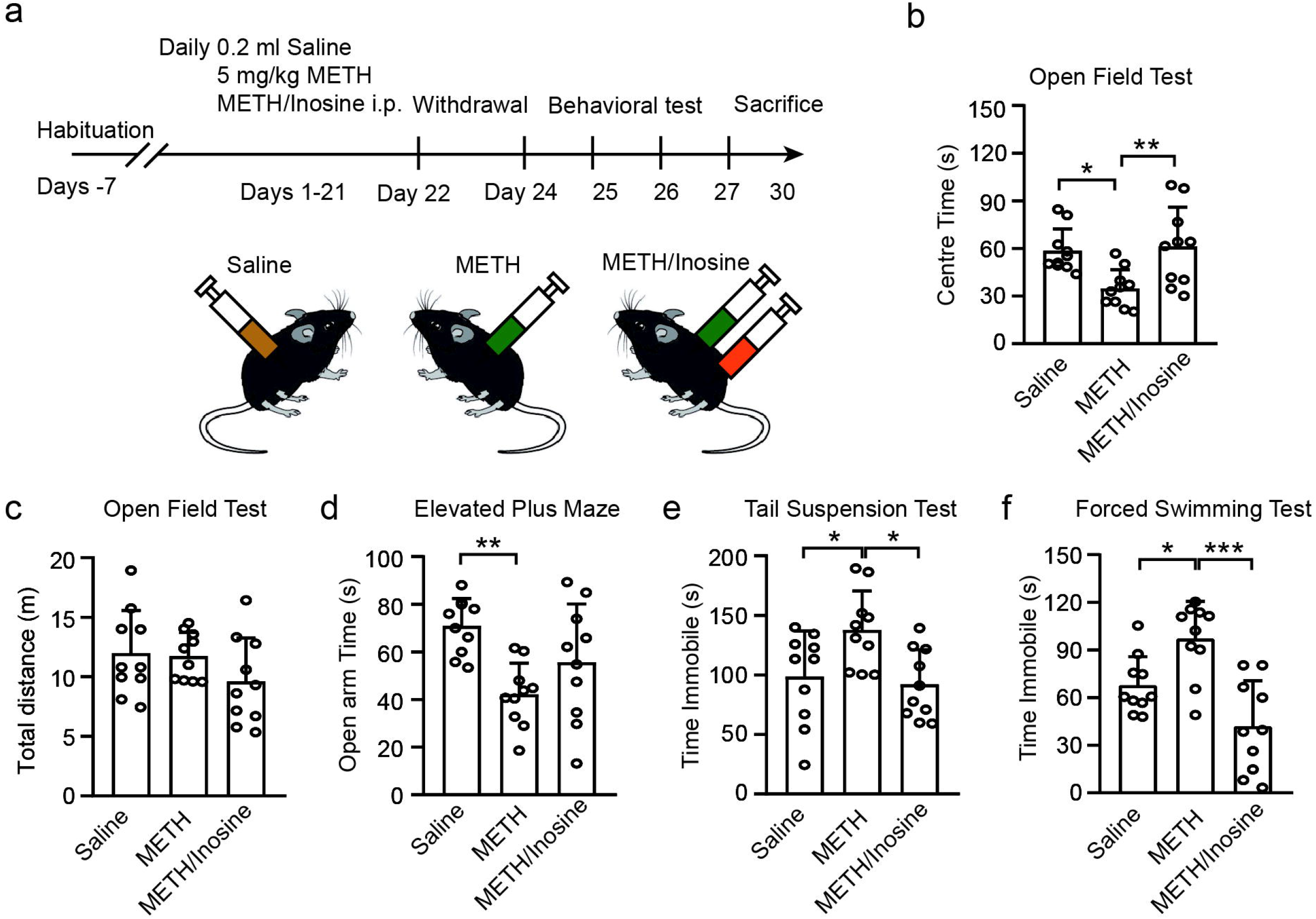
Administration of inosine reduces METH-induced anxiety and depression related behaviours in C57BL/6 mice. (a) Experimental protocol. (b) Pooled data for time spent in the centre and (c) total distance in the OFT, (d) the time in the open arm in the EPM, (e) immobile time in the TST, and (f) FST (n=10/group).

## DISCUSSION

In the present study, using state-of-the-art integrative multi-omics technologies and FMT models, we provided evidence showing that 1) METH withdrawal-induced anxiety and depression-like behaviours are convertible and transmissible via gut microbiota in a mouse model and 2) the therapeutic effects of metformin on psychiatric manifestations are associated with microbiota derived metabolites. Our results highlight the role of gut microbiota as an important mediating factor in substance withdrawal symptoms through the microbiota-gut-brain axis and its impact on host metabolism.

Cumulative evidence has shown that the differential pattern of gut microbiota could identify patients with various psychiatric or neurodevelopmental diseases and mediate relevant behavioural deficits [42, 43, 44]. Emerging studies reported that the exposure and the cessation of METH or other substance induced behavioural disturbances as well as alterations in gut microbiota [38, 45, 46]. Although these compelling association studies in humans suggest gut microbiota may impact psychiatric symptoms, a direct contribution by the microbiota to the pathophysiology and behavioural outcomes during METH withdrawal stage has not been well described. By analyzing human MAs and METH dependent mice, we showed that both MAs and the mouse model exhibited serious withdrawal symptoms especially anxiety-and depression-related behavioural deficits (table 1, figure 2). Meanwhile, the gut microbiota from human MAs exhibited higher community diversity and distinct microbial structures comparing to those of HCs, which is opposite to the findings in many other neuropsychiatric diseases, such as Alzheimer’s disease [47], schizophrenia [48], schizophrenia [49], and autism spectrum disorders [50]. The result was somewhat unexpected. We suspect that the mechanism of METH-induced psychiatric symptoms might be different than in other mental and neurological diseases, and this might be related to symptoms such as hyperactivity and excitement in MAs. Notably, the abundance of the core microflora *Rikenellaceae* was positively correlated with the severity of anxiety and depression in MAs. Although a strong presence of *Rikenellaceae* has been reported in Alzheimer’s disease [47] and schizophrenia [48], according to our knowledge, this is the first study reporting the association of the relative abundance of *Rikenellaceae* and the severity of METH-induced withdrawal symptoms.

In addition to the association analysis, the most exciting part of our study is that we performed two types of FMTs to investigate the role of microbiota in development of behavioural deficits (figure 2,3). In the human-to-mouse FMTs, we observed that the withdrawal symptoms including anxiety and depression could be transferred from MAs to the GF mouse model. Subsequently, in the mouse-to-mouse FMTs, by comparing the recipients GF mice from METH-treated mouse donors vs. saline-treated mouse donors, we found that the FMT-METH mouse recipients exhibited apparent anxiety and depression-like behavioural deficits, confirming the critical role of gut microbiota in the pathophysiology of the substance withdrawal syndrome.

Furthermore, it is well known that metformin is mostly used in diabetes treatment. The beneficial effect of metformin on substance withdrawal-related symptoms has been recently reported, but the mechanistic study was mostly focused on the CNS [17]. Although an alteration of both gut microbiota and its metabolomics in response to metformin treatment may be the key for the interpretation of physiological outcomes, to date, no study has focused on the role of metformin on microbiota in MAs, and the molecular mechanisms of metformin in diseases other than diabetes are not fully deciphered. Toward this end, we carried out two series of mouse experiments to validate the function and to explore the possible mechanism. In one study, we administrated metformin in the METH-dependent mouse model and validated the therapeutic effects of metformin on METH-induced behavioural phenotypes. In another study, we conducted mouse-to-mouse FMTs and consequently compared the behavioural outcomes of recipient GF mice with METH-, saline- or METH/metformin-treated mouse donors. Our analysis clearly demonstrated that the addition of metformin ameliorates METH withdrawal-related anxiety- and depression-like behavioural disturbances in mice, which is consistent with previous study in rat. Intrigued by the abovementioned data, however, we conclude that the gut microbiota could act as an alternative route for metformin functioning with respect to psychiatric symptoms (figure 3,4).

Extending the analysis to the gut microbiota composition and function alterations in response to METH exposure and metformin treatment, we characterized the bacterial taxonomic composition of mouse fecal samples from before and after FMTs relative to their controls (figure 5a, c). Comprehensive investigation of bacterial families suggests that higher level of *Rikenellaceae* which were positively associated with anxiety and depression were observed both in humans and mice (figure 7a, c). Post FMT, the *Rikenellaceae* in the GF mouse model were similar to those in donor samples (figure 7b, d) indicating successful colonisation using the FMT protocol. Strikingly, metformin was able to decrease the level of *Rikenellaceae* and ameliorates METH induced anxiety and depression-like behaviours in the recipient GF mice (figure 7b, d), suggesting that relative abundance of *Rikenellaceae* in gut microbiota could be used as a diagnostic biomarker for METH withdrawal syndrome.

**Figure 7.**
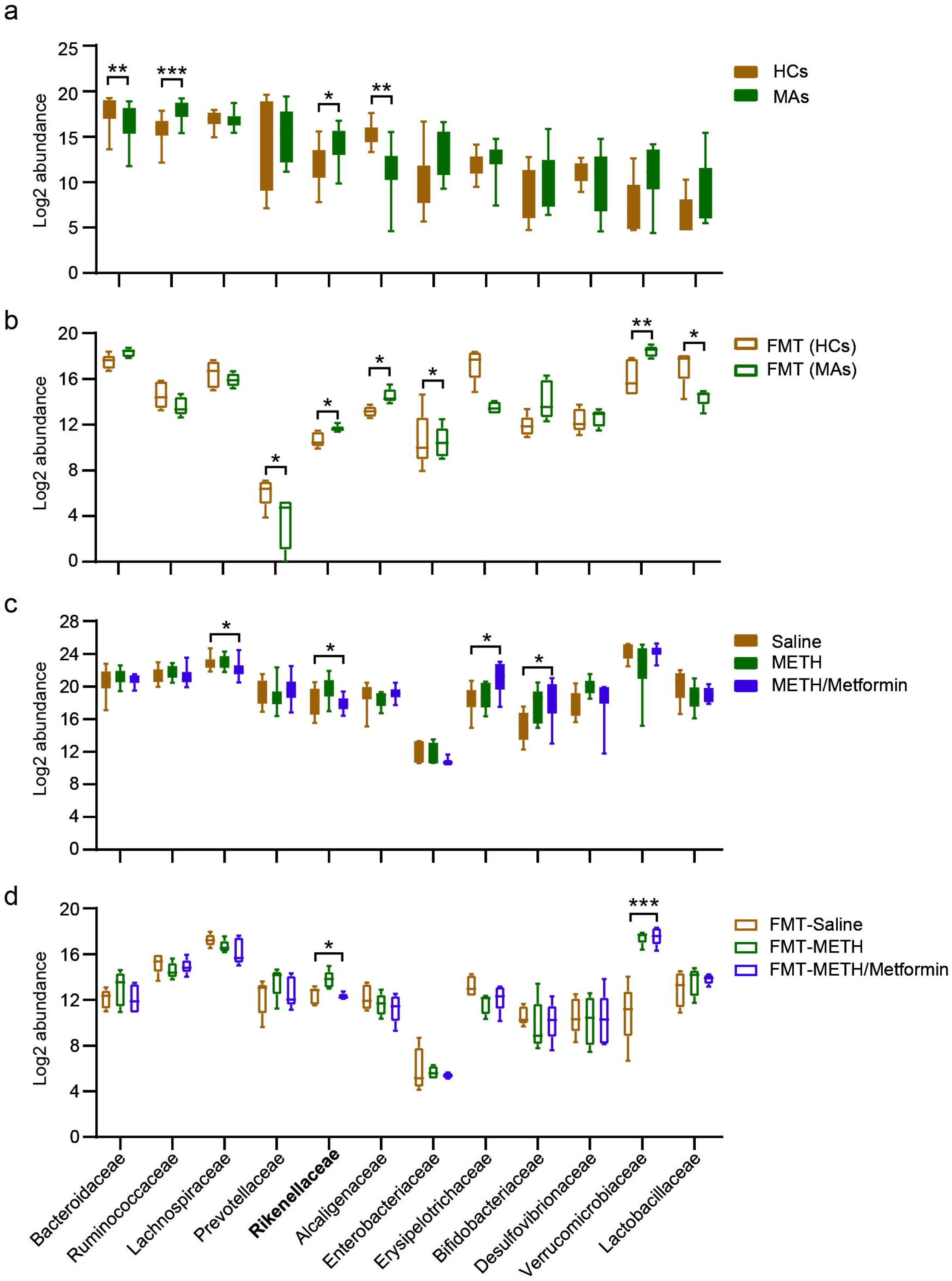
An overview of the fecal microbiota of MAs subjects, donor FMT mice or recipient FMT mice for the relative abundance of bacterial families. (a) Log2 abundance of the gut microbiota in MAs (n = 15) and HCs (n = 17) at the family level. (b) Comparison of intestinal microbiota at the family level in FMT-MAs mice (n = 5) and FMT-HCs mice (n = 5). (c-d) Comparison of abundant families among donor mice (n = 10/group) and post-FMT recipient mice (n = 5/group). A Wilcoxon-test was performed to determine if there were significant differences.

Simultaneously, we hypothesized that metabolites produced by gut bacteria might be an additional factor mediating psychiatric symptoms. The metabolic profiles calculated by PLS-DA were clearly distinct among saline, METH, METH/metformin mouse groups and in their corresponding FMT recipient mouse groups (figure 5a-b). Among identified differential metabolites, we discovered that as a key bacterial-derived metabolite inosine was reduced in fecal by METH exposure and could be restored by metformin treatment (figure 6a, c). Inosine, a major breakdown product of adenosine, has recently been shown to exert immunomodulatory [37, 51] and neuroprotective effects [41, 52] via the microbiota–inosine–A2A receptor axis or ERK and CREB pathway. In addition, inosine can permeate the blood–brain barrier and it is likely that decreased levels may reflect a reduction of these metabolites in the CNS. Oral administration of inosine has the potential to prevent depressive disorder [41]. Because of the low abundance in fecal samples of humans and mice, the intestinal microflora such as *B.pseudolongum* [37] and *L.reuteri* [51] that may generate amounts of purine/inosine have not yet been identified. There is no direct evidence that *Rikenellaceae* could regulate inosine production. Based on the evidence in the literature, we postulate that: 1) *Rikenellaceae* may promote inosine absorption by modulating the overall gut microbial community and structure; 2) metformin increases intestinal levels of short-chain fatty acids and activation of AMP-activated protein kinase and strengthens the intestinal barrier integrity to reduce inosine leakage, therefore resulting in the modulation of the *Rikenellaceae* level.

However, there are still some limitations in our study. One is the sample size of human MAs, and we need to obtain detailed information on the structure and function of the gut microbiota with a larger cohort study design. Second, 16S rRNA gene sequencing achieves only approximately 80% accuracy at the family level and is not able to fully resolve taxonomic profiles at the species or strain level [53, 54]. Therefore, shotgun sequencing will be warranted for the comprehensive profiling of the DNA from gut microbiota and to evaluate the effects and to unveil the subtle changes occurring with substance use and pharmacotherapy both in preclinical animal models and in randomised clinical trials.

In summary, our data provide support for the notion that the gut microbiota mediate some of the METH withdrawal-induced psychiatric symptoms, and metformin ameliorates behavioural deficits through an alternative route. Our metagenomic and metabolomic analyses reveal that the family *Rikenellaceae* and the metabolite inosine are the major mediators and contributors to the functional changes associated with METH use and metformin treatment. Overall, these results highlight the role of the gut microbiota in substance use disorders and the pathophysiology of withdrawal symptoms.

## Data availability

16S rRNA gene sequencing data for all samples have been submitted to the National Genomics Data Center (NGDC), under accession identification PRJCA001536 (http://bigd.big.ac.cn/gsa).

## Acknowledgements

This study was supported by the National Natural Science Foundation of China (No.3171101074, No.81870458, No. 31860306); Science and Technology Department of Yunnan Province (Grant No. 2018NS0086, 2019FE001, 202001AS070004, 202001AV070010); The Major project of Yunnan Provincial Bureau of Education (2020J0161; 2021J0234).

## Author contributions

Designed the experiments: JY and KW. Recruited clinical participants and collected samples: JY, ZZ, LB, PX, FC, HW, YZ, MZ. Animal behaviors and mice experiment: JY, YZ, MC, JG, FC. Performed the fecal microbiota transplantation: YZ, XY, MC, HL. Analyzed the 16S rRNA and metabolomics data: JY, KW, ZX, YK, QP, YD. Drafted the manuscript: JY, ZZ. All authors contributed to revision of the paper.

---

**Figure S1** Construction of the METH addiction model. (a) Timeline of the experimental sequence of locomotor sensitization. (b-c) METH (5mg/kg) treatment increased the locomotor sensitization and CPP scores in mice indicating the success of mice model establishment. (n =10 each group).

**Figure S2** Generation of microbiota depletion mice as GF mice by cocktail antibiotic-treated and colonization rates post FMT in mice-to-mice. Mice were given water containing antibiotic cocktail Ampicillin (1g/L), Metronidazole (1g/L), Neomycin(1g/L), and Vancomycin (0.5g/L) in the drinking water for 14 consecutive days. (a) Evolution of body weight gain before antibiotic treated (baseline) and after 14 days of antibiotic-treated (ABX). (b-c) DNA concentration and total DNA relative to the 16S rRNA gene for total bacterial load (all bacteria) in Saline or ABX group. DNA concentration expressed as ng/μL. (d) α-diversity indexes between the two groups. (e) Venn diagrams demonstrating the distribution of the OTUs shared among donors and recipient mice in mice-to-mice FMT model showed that most of the bacteria were found approached donor levels post-FMT indicating successful colonization and established communities from the donor microbiota.

**Figure S3** Evolution of body weight gain during the drug administration period. (a) Saline or Metformin treatment did not significantly change the body weight, while METH as well as METH/Metformin declined significantly with age. (b) Initial body weight and final body weight in inosine supplement test suggested that mice in METH as well as METH/inosine group weighed significantly less compared. (n = 10 for each group)

**Figure S4** The composition of fecal microbiota in the recipient mice. (a) Cladogram representing taxa enriched in fecal microbiota community of the three groups detected by the LEfSe tool. (b) Differential bacterial taxonomy selected by LEfSe analysis with LDA score >□3 in fecal microbiota community of the three groups.

