## Supplementary figures and images for "Metformin modulates microbiota-derived inosine and ameliorates methamphetamine-induced anxiety and depression-like withdrawal symptoms in mice"

### Supplementary Figure 1

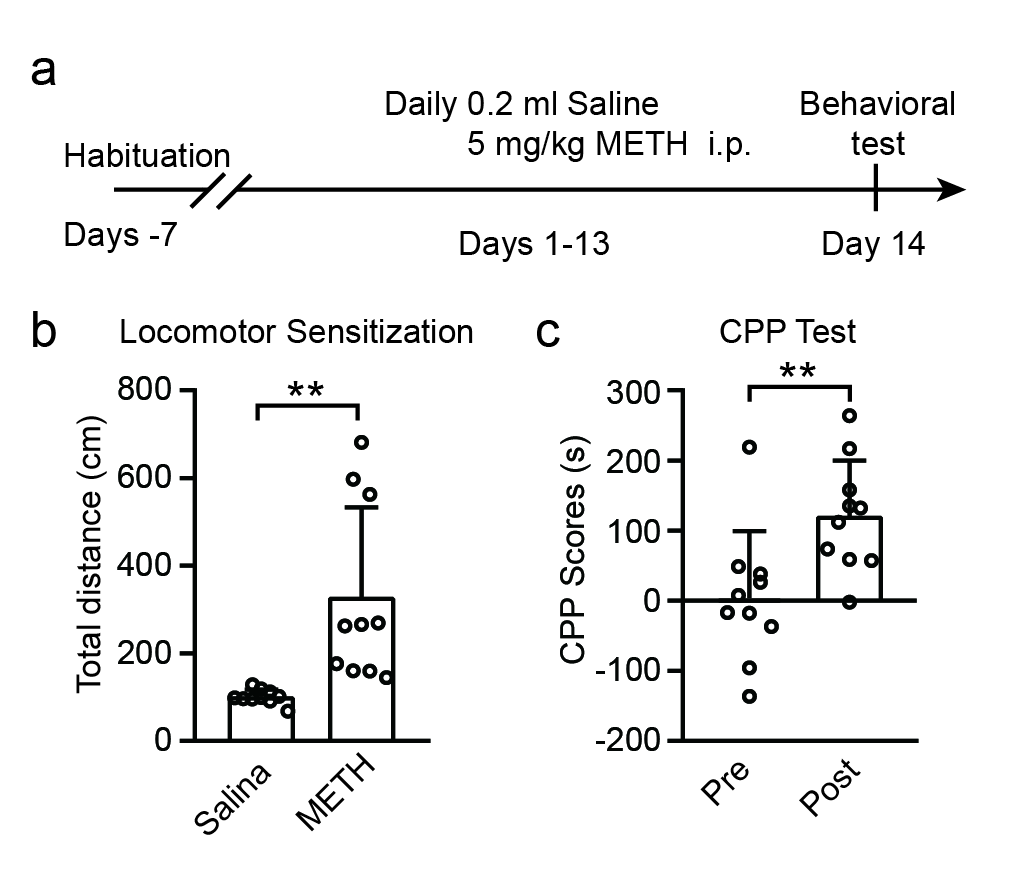

### Supplementary Figure 2

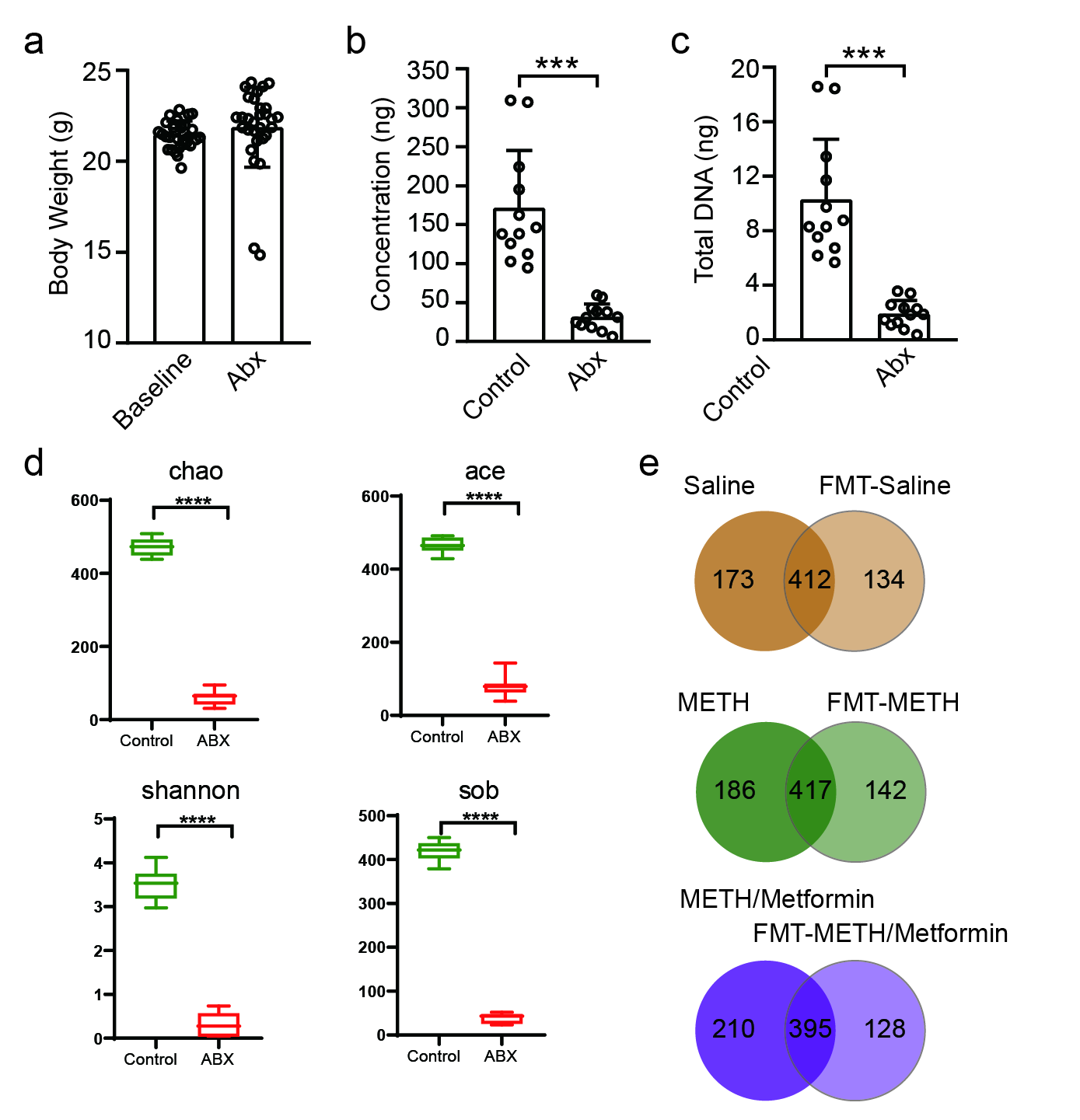

### Supplementary Figure 3

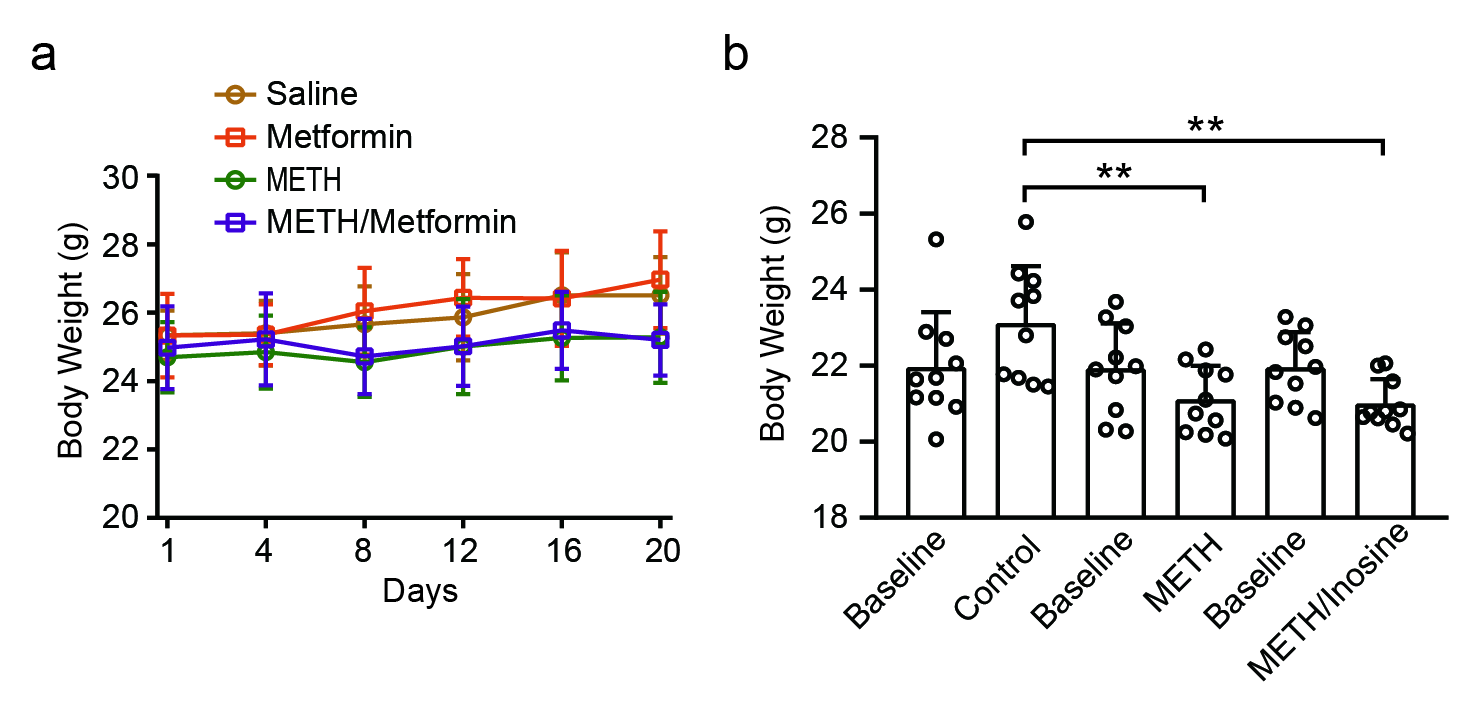

### Supplementary Figure 4

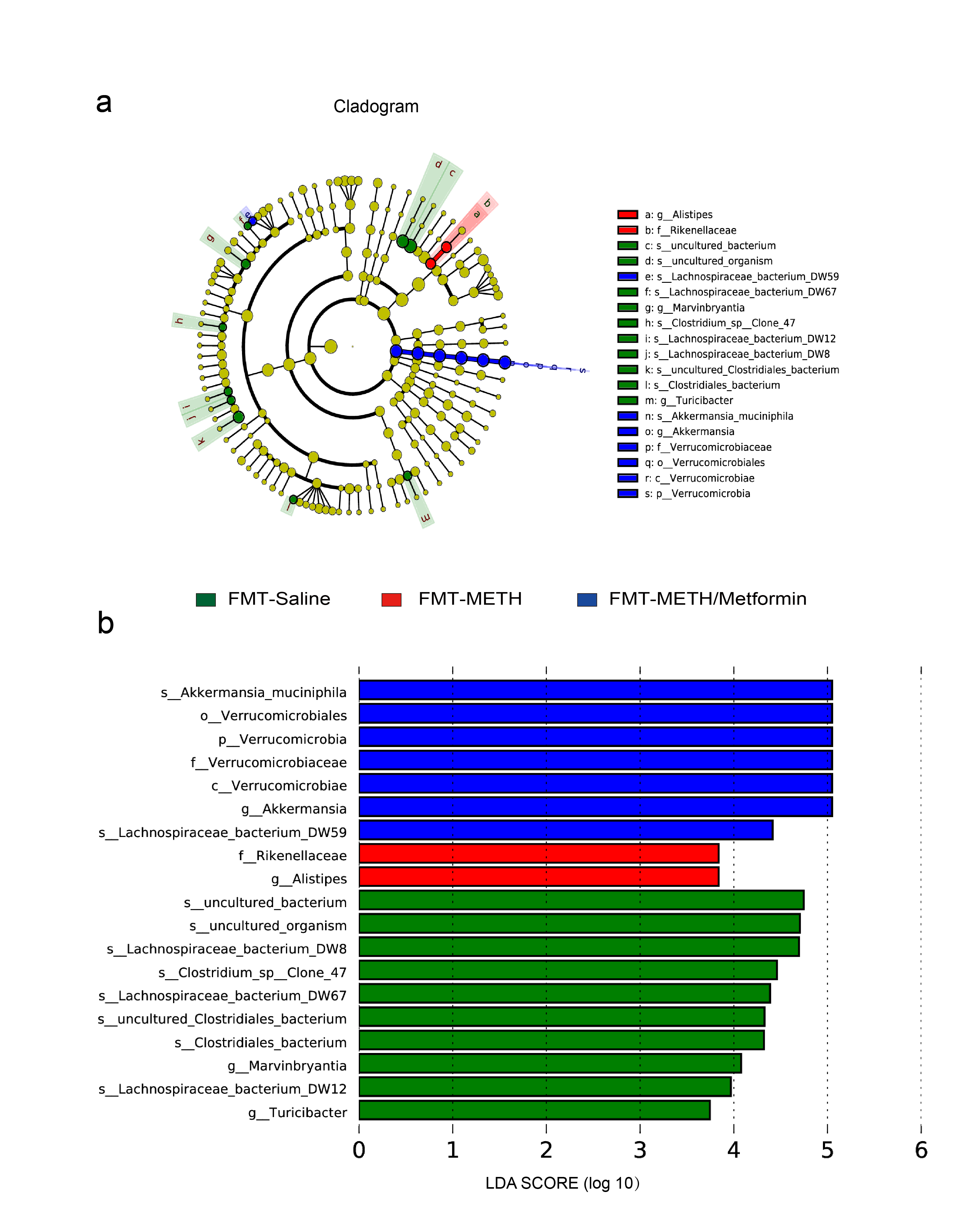
